# Gut microbial compositions mirror caste-specific diets in a major lineage of eusocial insects

**DOI:** 10.1101/418954

**Authors:** Saria Otani, Mariya Zhukova, N’golo Abdoulaye Koné, Rafael Rodrigues da Costa, Aram Mikaelyan, Panagiotis Sapountzis, Michael Poulsen

## Abstract

Eusocial insects owe their ecological success to the division of labour and processes within colonies often rely on the presence of specific microbial symbionts, but associations between microbial community compositions and castes with different tasks and diets within colonies remain largely unexplored. Fungus-growing termites evolved to use fungi to externally degrade plant material, complemented by specific and complex gut microbiotas. Here we explore to which extent division of labour and dietary differences within fungus-growing termite castes are linked to gut bacterial community structure. Using amplicon sequencing, we characterise community compositions in sterile (worker and soldier) and reproductive (queen and king) termites and combine this with gut enzyme, microscopy, and *in situ* analyses to further elucidate sterile caste-specific microbiota compositions. Gut bacterial communities are structured primarily according to termite caste and genus. In contrast to the observed rich and diverse sterile caste microbiotas, royal pair microbiotas are extremely skewed and dominated by few bacterial taxa, reflecting the specialised dietary intake and unique, reproduction-centred lifestyle of the queen and king.

## Introduction

Division of labour is a cornerstone in eusocial insects (ants, termites, and a number of wasps and bees). Eusocial insect communities are formed of sterile and reproductive castes, with members of the worker force often being specialised into morphological castes (workers and soldiers) performing temporally determined tasks (age-dependent polyethism) (Eggleton 2011, Ferguson-Gow *et al.*, 2014, Johnson 2010, Li *et al.*, 2015). Division of labour and the engagement of a range of symbiotic relationships have played key roles facilitating eusocial insect success (Batra and Batra 1979, Engel *et al.*, 2009, Lanan *et al.*, 2016, Sapountzis *et al.*, 2015). Work has shed light on the presence and role of gut microbial communities hosted by social insects (e.g., Anderson *et al.*, 2012, Brune and Dietrich 2015, Brune and Ohkuma 2011, Engel and Moran 2013), but our understanding of symbiont community structure across castes has been restricted to only a few studies (e.g., Benjamino and Graf 2016, Berlanga *et al.*, 2011, Hongoh *et al.*, 2006, Kapheim *et al.*, 2015, Li *et al.*, 2015, Poulsen *et al.*, 2014, Tarpy *et al.*, 2015).

Termites evolved from cockroaches 150 million years ago (MYA), and they exhibit the full range of sociality traits, from social structure more resembling cooperative brood care to strict division of labour with phenotypically-diverse colony members in the higher termites (Eggleton 2011). Microbial symbionts also play a pivotal role in all termites, primarily for food digestion and immunity (e.g., Brune 2014, Brune and Dietrich 2015, Douglas 2015). Lower and higher termite gut microbiotas are compositionally distinct, shaped by dietary differences and evolutionary histories between hosts and symbionts (Dietrich *et al.*, 2014, Donovan *et al.*, 2001, Eggleton and Tayasu 2001, Mikaelyan *et al.*, 2015a, 2017; Bourgoignon *et al.*, 2018). The higher termites diversified their diets 50 MYA to include soil, humus, leaf litter, grass, dung, and fungal material (Donovan *et al.*, 2001), coinciding with the loss of the lower termite-associated gut protists (Rahman *et al.*, 2015, Brune and Ohkuma 2011, Brune 2014) and diversification of primarily bacterial gut assemblies (Brune and Dietrich 2015).

Colony development is similar across higher termite lineages, with reproductive alates dispersing from mature colonies to pair up, found new colonies, and raise the first generation of workers (Eggleton 2011). When colonies mature, reproductive alates are produced (Eggleton 2011), and these likely bring the necessary gut microbes from their natal nest as an inoculum for the first worker cohort (Brune and Dietrich 2015, Rahman *et al.*, 2015, Benjamino and Graf 2016). Higher termite colonies contain both sterile (workers and soldiers) and reproductive castes (queen and king), the latter being monogamous for life (Boomsma 2009, Crespi and Yanega 1995). Some termite genera have a single sterile caste while others have dimorphic (major and minor) worker/soldier castes (Eggleton 2011, Korb and Hartfelder 2008). Reproductives are the only fertile individuals, while workers forage for food, maintain colony structures, and care for the brood and the royal pair, and soldiers defend the colony (Eggleton 2011).

The subfamily Macrotermitinae of higher termites exhibits further complexity through symbiotic association with an externally maintained fungal symbiont in the genus *Termitomyces* for nutritional and digestive functions (Sands 1969, Aanen *et al.*, 2002). In most Macrotermitinae species, the first workers emerging after colony foundation collect *Termitomyces* spores from the environment and culture these on a fungus comb consistent of young worker faecal deposits (Nobre *et al.*, 2011, Sands 1960). *Termitomyces* grows within these combs to produce nutrient- and conidia-rich nodules that are consumed by young workers (Badertscher *et al.*, 1983, Rouland-Lefèvre *et al.*, 2006), who redeposit conidia mixed with plant substrates as new comb. Old workers ingest the mature fungus comb (Badertscher *et al.*, 1983, Leuthold *et al.*, 1989) and the soldiers, queen, and king are fed fungal material by workers (Gerber *et al.*, 1988, Hongoh *et al.*, 2006, Leuthold *et al.*, 2004).

Obligate fungiculture in Macrotermitinae has altered the worker gut microbiota to be distinct from other termites and more similar to cockroach gut communities (Dietrich *et al.*, 2014, Leuthold *et al.*, 2004, Otani *et al.*, 2014). Qualitatively, individuals within a colony harbour a similar gut microbiota, likely due to within-colony transmission of gut bacteria (Hongoh 2010, 2011), but quantitative differences are poorly resolved. In *Macrotermes gilvus*, Hongoh *et al.*, (2006) showed that worker and soldier gut communities were more associated with termite age than caste. More recently, Poulsen *et al.*, (2014) found comparable microbiotas of workers and soldiers in a single *Macrotermes natalensis* colony, based on genus-level bacterial classification. Lastly, Li *et al.*, (2016) found variability in gut communities between different worker gut compartments and ages in *Odontotermes formosanus*. We thus lack a fundamental understanding of microbiota variability between castes from different fungus-growing termite species and within and between colonies from geographically distant sites.

We hypothesised that the presence of strict division of labour and resulting distinct dietary intakes of morphological castes should shape gut microbial composition in the Macrotermitinae. To address this, we performed gut microbiota composition analyses using MiSeq amplicon sequencing of workers and soldiers from three termite species in South Africa from colonies spanning over 350 km. We further applied a combination of fluorescence *in situ* hybridisation (FISH), light and confocal microscopy, and enzymatic profiling to elucidate *in situ* bacterial location and enzymatic differences across worker and soldier castes. We also, for the first time, perform comparative analyses of queen and king gut microbiotas from eight species of fungus-growing termites from the Ivory Coast to explore whether the strict fungal diet and the reproductive function shape gut microbial community compositions.

## Materials and Methods

### Termite samples and DNA extraction

Minor and major workers and soldiers from five colonies of *Macrotermes natalensis*, five colonies of *Odontotermes* sp., and one colony of *Odontotermes* cf. *badius* were collected in South Africa (Supplementary Table S1). At least 50 individuals per caste per colony were collected aseptically and stored frozen in RNA*later*^®^ (Ambion Inc., Invitrogen, CA) until DNA extraction. For reproductive castes, a single fungus-growing termite queen and/or king was collected from seven different *M*. *natalensis* colonies and two *Macrotermes* sp. colonies in South Africa and one colony per species of *Ancistrotermes cavithorax, Ancistrotermes guineensis, Macrotermes bellicosus, Odontotermes* sp., and *Pseudacanthotermes militaris* from the Lamto reserve in central Ivory Coast. Due to a lack of previous termite royal pair microbiota analyses, a grass-feeding higher termite queen from *Trinervitermes geminatus* from Ivory Coast was included for comparison. Ten whole guts were dissected and pooled from each sterile caste, while a single queen or king gut was used as a reproductive caste sample. DNA extractions were performed in triplicate using the DNeasy Blood and Tissue Kit (Qiagen, Hilden, Germany) following the manufacturer’s instructions, and DNA yields were assessed spectrophotometrically using NanoDrop ND-1000 (Thermo Fisher Scientific, USA).

### MiSeq amplicon sequencing

The V4 region of the 16S rRNA gene was amplified using v4.SA504 and v4.SB711 primers (Otani *et al.*, 2016) and dual-indexing sequencing strategy (Kozich *et al.*, 2013). PCR and library preparations were performed as in Otani *et al.*, (2016), and the SequalPrep Normalization Plate Kit (Life Technologies, Invitrogen) was used for library normalization. Sample concentrations were measured using the Library Quantification kit for Illumina platforms (Kapa Biosystems, Wilmington, MA). The library amplicon size was determined using the Agilent Bioanalyzer High Sensitivity DNA analysis kit (Agilent Technologies, CA). Samples were sequenced on the Illumina MiSeq platform using the MiSeq Reagent Kit V2 500 cycles (Illumina) (Koenigsknecht *et al.*, 2014, Kozich *et al.*, 2013). Mothur v. 1.34.3 (Schloss *et al.*, 2009) was used for analyses using the standard operating procedure (SOP) (http://www.mothur.org/wiki/MiSeq_SOP; Kozich *et al.*, 2013). Filtered sequences were aligned against the manually curated database DictDb 3.0 (Mikaelyan *et al.*, 2015b). Alignments were assigned to taxa with a confidence threshold of 80%, and operational taxonomic units (OTUs) were calculated at 3% species level classification. To compare alpha and beta diversity among groups, we performed a second round of analyses on the combined samples using essentially the same settings in mothur. Finally, dada2 was used to analyse the two datasets (sterile and royal) separate and merged, following the SOP https://benjjneb.github.io/dada2/tutorial.html page. This allowed us to use an alternative approach to analyse the data with a different output (Amplicon Sequence Variants which are similar to unique sequences in mothur) and a more aggressive filtering approach. Rarefaction curves using the 97% sequence similarity-generated OTUs (or the unique ASVs from the dada2 analysis) were constructed in R (R Core Team 2013).

### Analyses of caste gut microbiota composition

Two ordination analyses based on Bray-Curtis distances were performed in R to determine compositional similarities between sterile caste gut communities and between royal pair gut communities and visualised using PCoA (Principal Coordinates Analysis) and hierarchical clustering dendrograms, respectively. Analysis of molecular variance (AMOVA) was performed using R scripts implemented in Mothur (R Core Team 2013) to test whether distantly clustered microbial communities were significantly different (Excoffier *et al.*, 1992). To visualise the OTUs that contributed the most to gut community differences, bacterial taxa that based on PCoA loading values explained most of the dissimilarities between microbiotas, and that cumulatively accounted for >70% of the mean total abundance per community, were plotted as ternary plots in R. Then, their relative distributions in each caste were calculated and visualised in R. To allow for a phylogenetic comparison of the bacteria dominating royal pair guts, we performed a maximum likelihood (ML) analysis on the 45 most abundant bacterial taxa (collectively accounting for >0.1% summed abundance across all samples) using the Tamura-Nei model in MEGA and FigTree (Tamura *et al.*, 2007) visualised in R. Shannon and Evenness indices were calculated using an R script implemented in mothur. Each alpha-diversity index was fitted in a linear mixed model, with caste, genus, and site as explanatory variables and colony as a random variable (nlme package in R; Pinhero *et al.*, 2018). Multiple comparisons among groups were performed using Tukey post-hoc tests in R. For beta-diversity comparisons we used the betapart package in R, which partitions beta-diversity into turnover and nestedness (Baselga and Orme 2012). Similar to the alpha-diversity calculations, we used the merged dataset to test for the effects of caste, genus, and site on overall diversity, turnover, and nestedness. Multiple comparisons were also performed using Tukey post-hoc tests implemented in the betapart package. To examine the differential representation of OTUs among castes, we inserted the unrarefied merged OTU tables of mothur and dada2 in the DAtest package in R (Russel *et al.*, 2018), which suggested the use of DESeq2 (Love *et al.*, 2014).

### Light microscopy

Three termites from each sterile cast of *Odontotermes* cf. *badius* (Od152), *Odontotermes* sp. (Od127) and *Macrotermes natalensis* (Mn173, Mn160) were dissected in 0.01M phosphate buffer (pH 7.4), and their guts were fixed in 2.5% glutaraldehyde (Sigma) in 0.1 M sodium cacodylate buffer (pH 7.4) for 2.5 h. This was followed by washings in the same buffer and post-fixation in 1% OsO_4_ for 1 h, after which samples were placed in a 1% aqueous solution of uranyl acetate and left at 4°C overnight. Samples were then dehydrated in an ethanol series and acetone and embedded in Spurr low-viscosity resin (Ted Pella Inc.). Sections of 500 nm thick were stained with toluidine blue and observed under a light microscope (Nikon Eclipse 80i).

### Fluorescence in situ hybridisation (FISH) and confocal microscopy

Three termites from each sterile caste of *O.* cf. *badius* (colony Od152) and *Macrotermes natalensis* (colony Mn173) were dissected in phosphate buffered saline (PBS) and their guts were placed in 4% paraformaldehyde for at least 24 h. Samples were washed with distilled water and stored in 70% ethanol at 4°C before performing the following steps. Fixed guts were dehydrated via graded alcohol and Histoclear (Sigma-Aldrich, St Louis, MO) and embedded in Paraplast Plus (Sigma-Aldrich). After cutting 6 to 8 µm sections and mounting them onto glass slides (TruBond380, Tru Scientific, Bellingham, WA), sections were deparaffinised in methylcyclohexane and air-dried. An extra lysozyme (1 mg/ml) in 0.01 M Tris-HCl buffer (pH 8), 0.005M EDTA treatment was performed at 37°C for 5 min to permeabilise Gram-positive bacteria (Firmicutes) followed by washing with PBS. Samples were then incubated with 0.1 mg/ml pepsin in 0.01N hydrochloric acid for 10 min at 37°C and washed with PBS. For hybridisation, we used the protocol described previously (Koga *et al.*, 2009). All probe sequences are listed in Supplementary Table S7. The affinities probes were checked using the R “compute.affinities” function (R Core Team 2013). Universal bacterial probe EUB388 was used simultaneously with specific probes as a positive control, and the reverse EUB388 (nonEUB388) probe was used as a negative control (Supplementary Table S7; Supplementary Figure S4). No bacterial signal was detected with the negative probes (Supplementary Figure S4). Samples were incubated at room temperature for hybridisation with all probes except SPIRO1400, which was hybridised at 35°C. After being thoroughly washed with PBS-TX buffer (PBS, 0.3% TritonX-100), samples were mounted in Vectashield medium with DAPI (Vector Laboratories Ltd., Peterborough, UK) and observed by confocal microscopy (LSM 700, Zeiss).

### Enzyme activities of sterile caste gut communities

To determine differences in gut enzyme capacities across different termite castes, we used sterile caste samples from six colonies of *M. natalensis*, three colonies of *Odontotermes* sp., and three colonies of *O.* cf. *badius*. Seventeen azurine-crosslinked (AZCL) substrates were used, and screening media were prepared using 0.1g/l of AZCL substrates in agarose (1% agarose, 23 mM phosphoric acid, 23 mM acetic acid, 23 mM boric acid). The pH was adjusted for each substrate according to the manufacturer’s instructions (Megazyme, Bray, Ireland). Briefly, 100 mg of each sample (ca. 20 termite guts) was crushed with a pestle in a 1.5 Eppendorf tube containing 1000 µl distilled water followed by vortexing and centrifugation (15 000 x g). Supernatants were retrieved and applied to AZCL assay plate wells (ca. 0.1 cm^2^) in triplicate technical replicates. After 22-24 h incubation at 25°C, plates were photographed, and enzyme activity quantified by measuring the halo areas around the wells using ImageJ ver. 1.6.0. A PCoA analysis based on Bray-Curtis distances was performed in R (R core team). To test whether PCoA clustering of gut enzymatic capacities were significant, one-way PERMANOVA analysis was performed between caste clusters using PAST ver. 2.17c (Hammer *et al.*, 2001).

## Results and Discussion

### Sequencing and bacterial identification

Sequencing the bacterial communities of 180 sterile and reproductive termite caste gut samples resulted in 3 047 202 quality-filtered sequences and 12 610 bacterial OTUs (Supplementary Tables S1-S3). Rarefaction analyses indicated that sequence coverage was sufficient to capture community diversity, with the reproductive castes having a ten-fold lower diversity than sterile castes (Supplementary Figure S1). Family-level classification success using the DictDb 3.0 database (Mikaelyan *et al.*, 2015b) was 74% and 84% in sterile and reproductive castes, respectively (Supplementary Tables S2-S3). mothur produced 9 314 OTUs across worker and soldier guts from 33 phyla, dominated by taxa previously reported to be abundant in fungus-growing termites (Dietrich *et al.*, 2014a, Mikaelyan *et al.*, 2015a, Otani *et al.*, 2014). The dada2 analyses were largely consistent with our mothur analyses (Supplementary Tables S8, S9), but we nevertheless performed alpha- and beta-diversity analyses on both classification approaches to support our main conclusions.

### Compositional and functional differences across sterile caste gut communities

The vast majority of the identified taxa were evenly distributed across minor workers, major workers, and soldiers in *M. natalensis* and the two *Odontotermes* species (centres of the ternary plots, Figure 1A). This included some of the most abundant bacterial OTUs in the genera *Alistipes* and *Treponema*, which is likely driven by within-colony transfers through mouth-to-mouth (stomodeal) or anus-to-mouth (proctodeal) trophallaxis (Hongoh 2010, Nalepa 2015). However, some bacteria (Figures 1, 2 and Table S12) were differentially-abundant across castes, evident from both ternary plots and subsequent DESeq2 analyses (Table S12). The latter showed that the majority of OTUs that varied between castes produced significant contrasts in at least one pairwise DESeq2 comparison (Figure 1, Supplementary Table S12). We cannot rule out that more may be differentially abundant, as our cut-off of 70% cumulative abundance would select against relatively rare OTUs that could differ in abundance between castes. The significant differences were almost exclusively driven by worker *versus* soldier comparisons and only rarely driven by differences between minors and majors (Table S12). *Arcobacter* and *Enterobacter* 4 were the only two OTUs that were common in workers of both termite genera (according to Figure 1B) and similarly the DESeq2 analysis showed that the vast majority of the differentially distributed OTUs were present only in *M*. *natalensis* or *Odontotermes* spp. (Figure 1B; Table S12). These results did not change in any appreciable way in the ASV output table from the dada2 analysis (Supplementary Table S12). Phylogenetic analyses of these OTUs revealed that although they were not identical between termite genera, many were sister taxa, indicating potential diversification from common ancestors (Supplementary Figure S2). If so, this would imply that a distinct set of OTUs may show differential presence and represent ancient specialised roles within the guts of respective termite castes.

**Figure 1.**
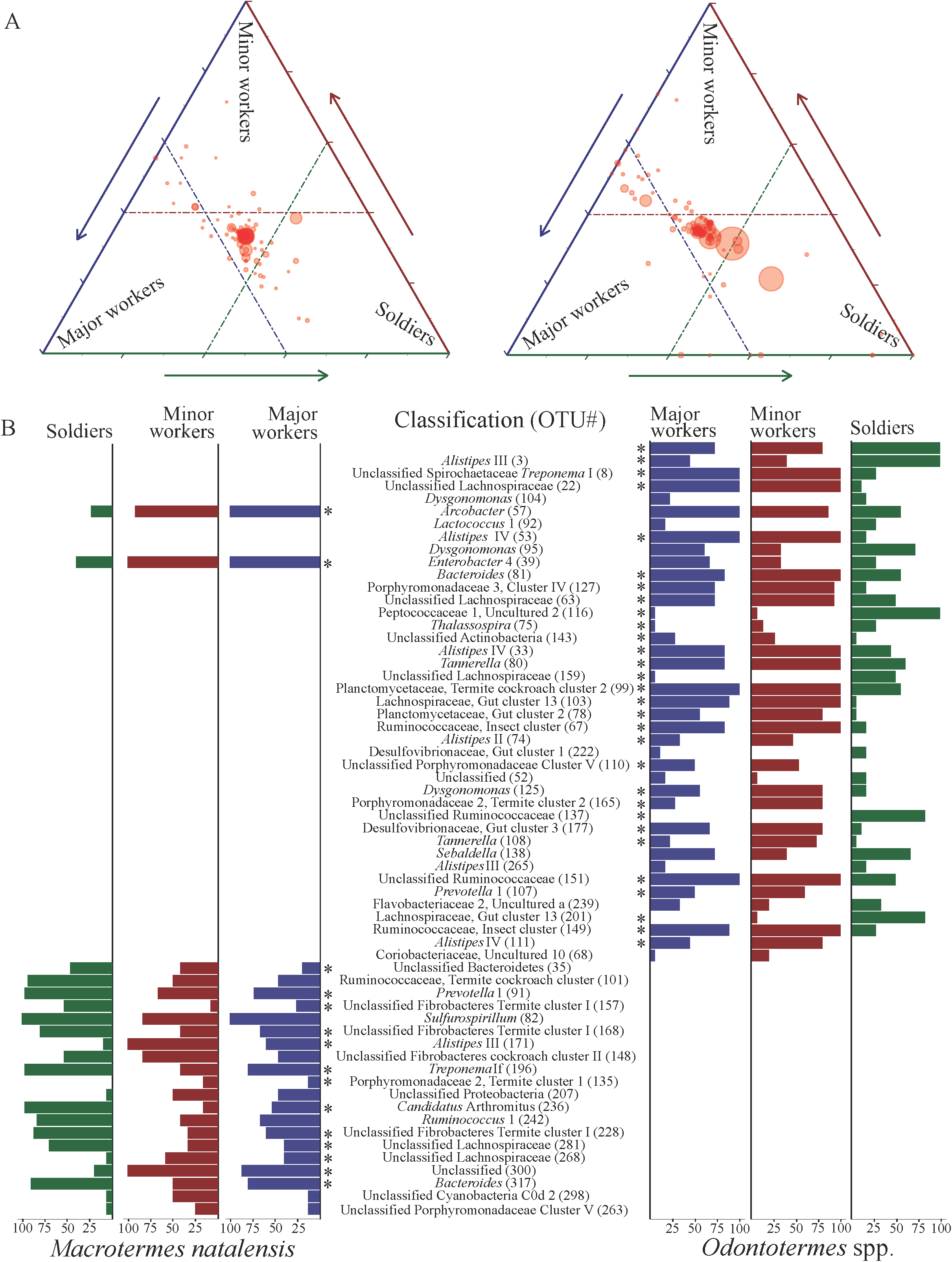
Distribution and differential abundance of gut bacteria across sterile castes. A) Ternary plot of the most abundant bacteria (>70% relative abundances across gut communities) and their distribution between sterile castes. Each circle represents an OTU, and its size represents the relative abundance. The position of a circle represents the bacterium contribution to differentiating caste microbiota compositions, where the dashed lines represent 40% of the dissimilarity explained in the ordination analyses (based on the PCoA loading values). B) The proportional distributions of bacterial OTUs within a caste that only explain >40% of the microbial community variations between castes in a termite genus. The scale represents the proportion of an OTU in a caste, i.e., 100 means it is present in all individuals of a given caste. OTUs significantly different in at least one pairwise DESeq2 comparison are indicated with an asterisk (for the full results, see Table S12).

**Figure 2.**
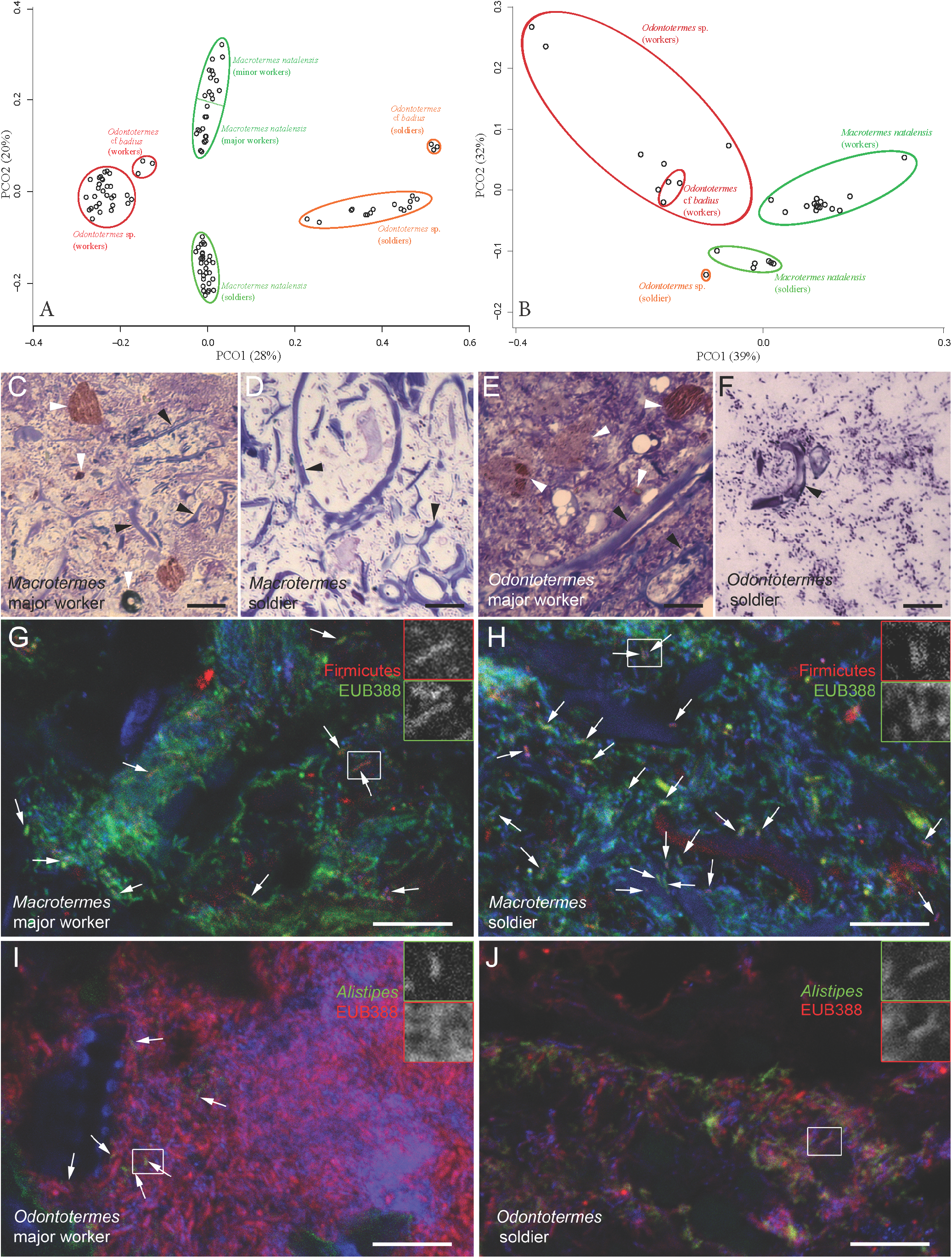
Structural and functional differences between gut communities of sterile castes. A) Bacterial community similarity analysis of the 108 sterile caste gut microbiota, visualised using principal coordinate analysis (PCoA) based on Bray-Curtis distances with each circle representing a gut microbiota. *p* values of Bray-Curtis distances between the eclipses were tested using an AMOVA test (*P* <0.001, Supplementary Table S6). B) PCoA of AZCL enzyme activities in different castes and ages. C-F) Different substrates in paunches of major workers and soldiers of *Macrotermes natalensis* (C, D) and *Odontotermes* cf. *badius* (E, F). Fungal material marked with black arrowheads, other types of substrate indicated by white arrowheads. Light microscopy of semi-thin sections stained with toluidine blue. G, H) representative images of fluorescent *in situ* hybridisation with *Firmicutes*-specific probe (red, arrows) in sections of the paunch of a *Macrotermes natalensis* major worker (G) and soldier (H). All bacteria were stained with the domain-specific probe EUB388 (green), DNA was stained with DAPI (blue). I, J) *Alistipes* III (green, arrows in I) in sections of paunch of an *O.* cf. *badius* major worker (I) and soldier (J), all bacteria were stained red (EUB388 probe), DNA was stained with DAPI (blue). *Alistipes* III was abundant in soldiers of *O.* cf. *badius* (J). The inserts show overlapping staining with two probes of target bacteria at higher magnification. Scale bars are 10 µm.

Differentially occurring bacteria are likely to represent OTUs that serve disparate functions in fungus-growing termite castes, supported by differences in enzymatic profiles in caste guts. The different enzymatic capabilities have to be explained by (a) differences in OTU composition or (b) differences in the termite enzymatic profiles, with the former being most plausible. Consistent with this assertion, *Alistipes* IV was more abundant in *Odontotermes* spp. workers than soldiers (Figure 1B), and carbohydrate-active enzyme analysis of *M. natalensis* guts have suggested that *Alistipes* lineages have the enzymatic capacity to hydrolyse chitin and other fungal cell wall components (Liu *et al.*, 2013, Poulsen *et al.*, 2014). While Lachnospiraceae (Firmicutes) are believed to be involved in non-cellulosic saccharide degradation in fungus-growing termites (Li *et al.*, 2016, Veivers *et al.*, 1991, Watanabe and Tokuda 2010), consistent with a number of Lachnospiraceae OTUs being mainly present in soldiers (Supplementary Table S2; Figure 1B). *Treponema*, a metabolically versatile genus involved in lignocellulose breakdown in other termites (Droge *et al.*, 2008, Graber and Breznak 2004, Mikaelyan *et al.*, 2014) is generally scarce in fungus-growing termites (Rahman *et al.*, 2015, Dietrich *et al.*, 2014, Otani *et al.*, 2014), but was nevertheless differentially distributed between sterile castes (Table S12).

When using the two separate analyses (reproductive and sterile castes) the differences in communities were substantial enough to allow for distinct and significant separation of termite worker and soldier gut communities in ordination plots (Figure 2A; AMOVA *p* < 0.001; Supplementary Table S6), but the separate analyses did not allow for a comparison of reproductives with sterile castes directly. Within the sterile castes, the most distinct separations were between workers and soldiers, while separation within morphologically different castes (i.e., minor and major workers and minor and major soldiers, respectively) were less distinct (Figure 2A; Table S12). The effect of caste became even clearer when we used the merged datasets in a combined analysis and examined caste, genus, and site to show that the first two contribute the most to the observed differences, and where partitioning of the beta-diversity showed that species turnover (species replacement) was the main reason for these differences (Supplementary Table S11). This is consistent with recent work elucidating how the evolutionary histories of gut microbes associated with termites are shaped by mixed-mode transmission and acquisitions from environments (Bourgoignon *et al.*, 2018). The similarities within castes persist across geographical locations of colonies spanning more than 350 km (Figure 2A), consistent with previous studies showing that macrotermitine gut microbiotas are not only distinct from other termites but are also stable over geographical locations and time (Otani *et al.*, 2016, Otani *et al.*, 2014) and that differences between gut communities are less influenced by colony of origin (Hongoh *et al.*, 2006, Otani *et al.*, 2016). The significant contrasts for beta diversity did not change in any appreciable way when we used the merged dataset from the dada2 analysis.

### *Gut content characterisation of termite sterile castes and* in situ *detection of caste-specific bacteria*

Light microscopy revealed differences between worker and soldier gut substrates in the paunch, with fungal substrates dominating soldier guts and worker guts containing a mixture of soil particles, fungus and plant material (Figure 2C-F). These findings support the previously hypothesised predominant fungal diet of soldiers (Hongoh *et al.*, 2006, Leuthold *et al.*, 2004), while confirming that worker diets are more diverse (Eggleton 2011, Hongoh *et al.*, 2006). These differences imply the different nutritional/digestive requirements are likely supported by differences in gut microbiotas. Further support for this was provided by confocal microscopy and FISH analyses using probes specific for four bacterial OTUs in *Odontotermes* and one in *M*. *natalensis* (Supplementary Table S7), specifically Lachnospiraceae, Bacteroidetes, *Alistipes* III, and members of the Spirochaetaceae, including *Treponema* (Supplementary Table S12). The Lachnospiraceae OTU133 (Firmicutes) was more abundant and appeared more uniformly represented in *M*. *natalensis* soldiers compared to workers (Figure 2G, H), consistent with the sequencing analysis (Figure 1B, Supplementary Table S2, Table S12). This corroborates previous work showing that Lachnospiraceae are more abundant in *Odontotermes formosanus* young termites feeding mainly on fungal nodules or degrading non-cellulosic plant oligosaccharides (Li *et al.*, 2016) and contributing to lignin depolymerisation (Li *et al.*, 2017). Similarly, *Alistipes* III (OTU8, Figure 1B; Supplementary Table S2) was significantly more abundant in *Odontotermes* soldier guts compared to worker guts (Figure 2I, J), consistent with the sequencing results (Figure 1B, Supplementary Table S12). The remaining OTUs showed similar patterns, with morphologically different Bacteroidetes members being abundant in *Odontotermes* worker but not soldier guts (Supplementary Table S7; Supplementary Figure S3). This suggests that even though the Bacteroidetes phylum is abundant in all sterile caste guts, dietary variation between castes causes representative OTUs within the phylum to be differently abundant between worker and soldier guts, which in turn serves the division of labour by specializing different caste microbiotas to different catabolic functions. The probe targeting Spirochaetaceae was also more abundant in *Odontotermes* workers than soldiers (Supplementary Table S7; Supplementary Figure S3), further supporting the presence of caste-enriched bacterial communities. No bacterial signal was detected with the negative probes (Supplementary Figure S4).

### Worker and soldier guts differ in their enzymatic capacities

Bacterial composition variations between sterile caste communities were also apparent in their enzymatic capacities (Figure 2B). Worker and soldier guts from both *M. natalensis* and *Odontotermes* exhibited significant differences in their ability to degrade 14 different substrates (Figure 2B, Supplementary Table S4). *M. natalensis* workers showed significant enzymatic differences compared to their generic soldiers and *Odontotermes* sterile castes (Figure 2B, PERMANOVA *p* < 0.03), with the highest capacities to degrade all the substrates present in all castes (Supplementary Table S4). While *Odontotermes* worker guts showed significantly lower enzymatic capacities compared to *M. natalensis* worker guts (Figure 2B, PERMANOVA *p* < 0.02; Supplementary Table S4), *Odontotermes* soldiers did not show any significant differences compared to other castes (Figure 2B, PERMANOVA, *p* > 0.6; Supplementary Table S4). In both termite genera, worker guts showed a greater capacity to degrade a wider range of substrates and to degrade plant-derived substrates, e.g., HE-cellulose, xyloglucan, xylan and arabinoxylan, than soldiers (Supplementary Table S4), consistent with the more diverse diet of workers and supporting associations between the functional and compositional differences in caste guts.

### Queen and king gut microbiotas differ markedly from those of sterile termites

Previous compositional analyses of a single *M. natalensis* queen indicated that her gut was dominated by a single OTU, thought to be a *Bacillus* sp., determined by a low-resolution database for taxonomic binning (Poulsen *et al.*, 2014). Expanding on this finding, we found that one to three bacterial lineages dominate gut communities in each of twenty-three fungus-growing termite royal pairs (13 queens and 10 kings) and a queen from the grass-feeding termite species *T. geminatus*, resulting in ten bacterial lineages accounting for >80% of all sequences generated from royal pair libraries (Figure 3, Supplementary Table S3). Lactobacillales lineages dominated *M. natalensis, Macrotermes* sp., two *Ancistrotermes* species, and *P. militaris*, while the Desulfovibrionaceae gut cluster 1 OTU was the most abundant in a *M. bellicosus* queen gut (82.9% relative abundance) and *Sebaldella* dominated an *Odontotermes* sp. queen and king (53.6% and 82% relative abundance, respectively (Figure 3, Supplementary Table S3). Consistently, the dominant bacterial lineages in reproductives were absent or only present in low abundances in sterile caste guts (Supplementary Tables S2-S3), and OTUs that generally dominated sterile guts were absent or only present in trace amounts in royal pairs. The only exception to this was in queens of *M. natalensis* Mn118, Mn133, and Mn134 with more diverse and uniform bacterial communities (Figure 3).

**Figure 3.**
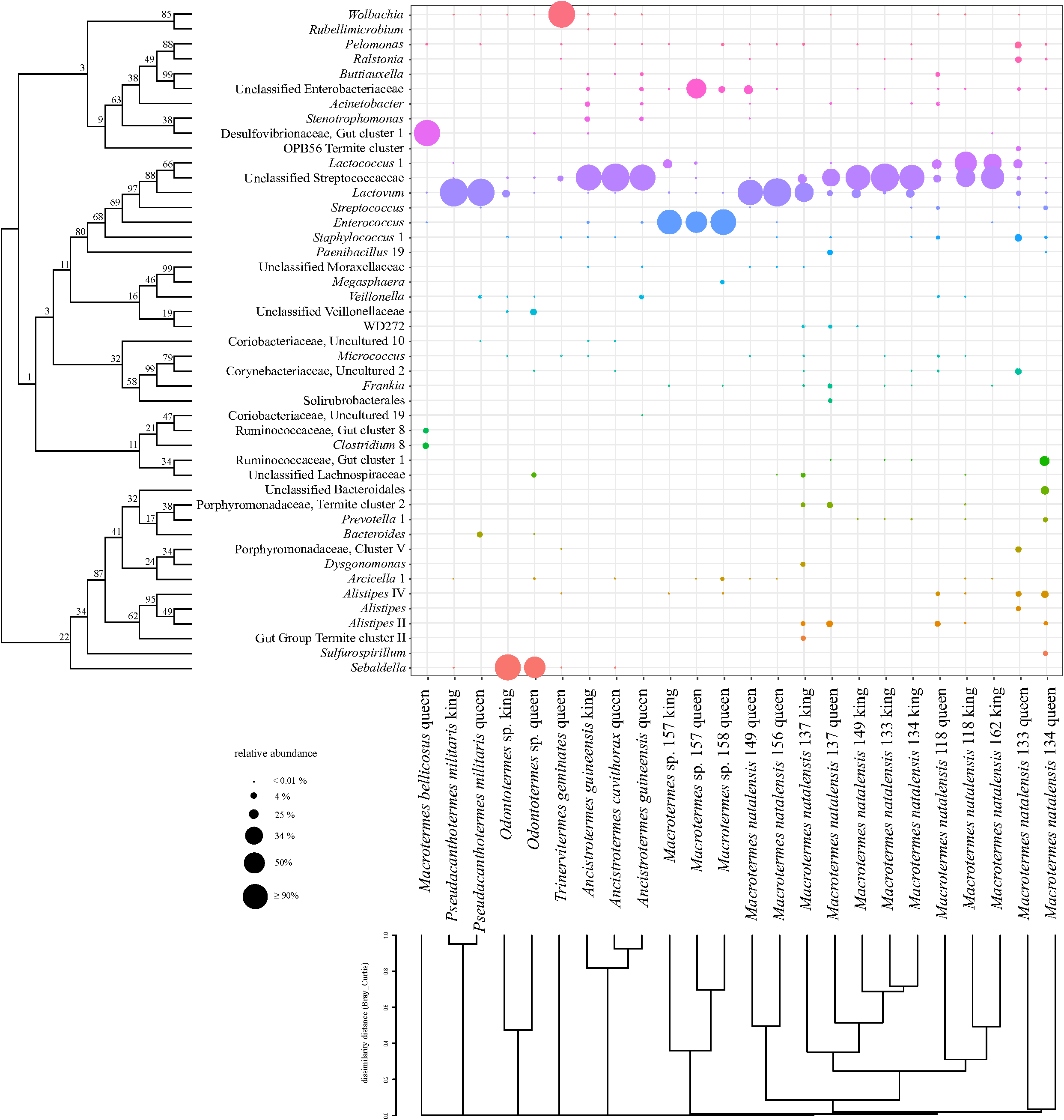
Gut bacterial communities in royal pairs from seven fungus-growing termite species and a grass-feeding *Trinervitermes geminatus* queen. The most abundant bacterial taxa in royal pair guts, accounting for >0.1% relative abundance in each community. Circle sizes indicate the relative abundance of a bacterium in the corresponding gut, and colours represent different bacterial OTUs, which totals >94% of all OTUs detected. *Wolbachia* dominated the grass-feeding *T. geminatus* queen gut only (Supplementary Table S3) driving this queen to be compositionally distinct from fungus-growing termite queen and king gut microbiotas (Supplementary Figure S5). The ML tree at the left represents the phylogenetic relationships between the bacteria and the dendrogram at the bottom visualises gut microbiota similarity analysis based on Bray-Curtis distances based on the most abundant bacterial taxa (dissimilarity analysis of the entire royal pair communities is presented in a PCoA plot in Supplementary Figure S5).

With few bacterial genera dominating queen and king guts, the communities were greatly skewed, in contrast to those of sterile castes (Figure 4), so that alpha-diversity of royal pair bacterial communities was significantly lower than in workers and soldiers (Figure 4, Supplementary Table S5). Neither sampling site nor termite genus significantly affected Shannon or evenness indices, while caste was highly significant for both mothur or dada2 analyses for both the separate and the merged datasets (Supplementary Table S5). This finding is in accordance with the highly significant contrasts in the comparison of beta-diversity of sterile and reproductive castes (Table S11), suggesting that caste is the most important factor shaping termite gut bacterial communities. There was also a strong association between community structure and termite host species, indicating that conspecific royal pairs more commonly associate with the same or closely related bacteria (Figure 3; Supplementary Figure S5; Supplementary Table S6).

**Figure 4.**
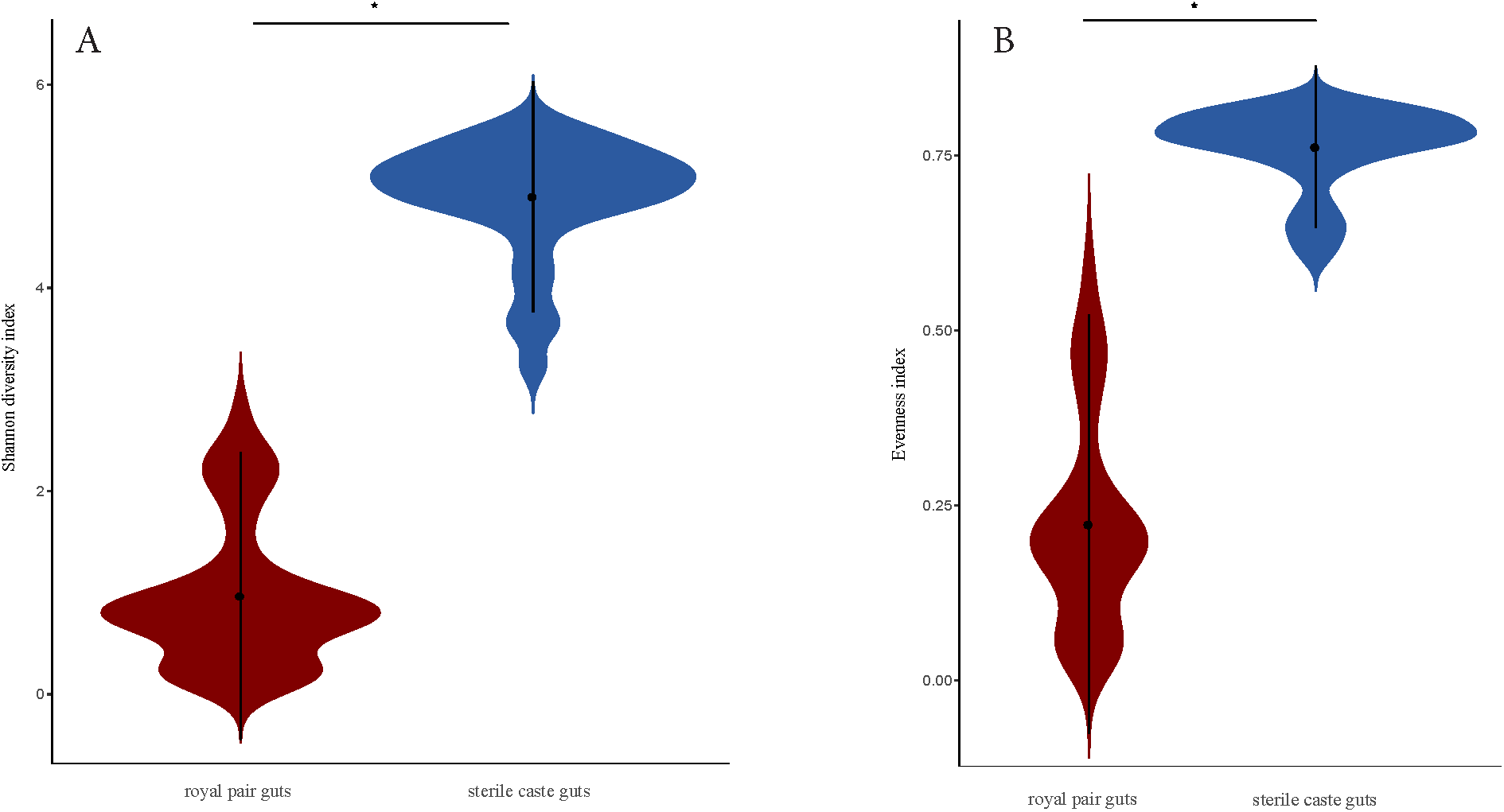
Diversity and evenness indices in gut microbiota from sterile and reproductive castes. A) Shannon diversity indices of sterile and reproductive caste gut communities calculated with an R-implemented script in Mothur and visualised by a violin plot in R. B) Evenness index of termite sterile and reproductive caste gut bacterial communities calculated with an R-implemented script in Mothur and visualised by a violin plot in R. * *p* < 0.00001 (*p*-values are from Mann– Whitney–Wilcoxon tests in R, Supplementary Table S5).

The presence of highly reduced diversity and royal pair specific microbiotas suggests that the strict fungal diet and reproduction-centred protected lifestyle, in which queens and kings are encapsulated in the royal chamber for the duration of their lifetime, have resulted in simplified gut microbiotas. This dramatic shift resembles the reported differences between workers and queens in the honeybee, where queens exhibit reduced microbial diversity and shifts in dominant gut bacteria (Kapheim *et al.*, 2015, Kwong and Moran 2016, Tarpy *et al.*, 2015). The dominant bacteria differ between honey bee and fungus-growing termite queens, but it is intriguing that the termite royal pair guts are dominated by members of the Lactobacillales, which appear to be involved in sugar uptake and fermentation in bee worker guts, and some lineages have antimicrobial properties (Ellegaard *et al.*, 2015, Killer *et al.*, 2014, Kwong *et al.*, 2014, Kwong and Moran 2016). It is conceivable that these termite royal pair gut microbes play similar roles but work to elucidate this is needed.

## Conclusions

To date, this is the most comprehensive survey of caste-specific microbiotas in fungus-farming termites, confirming predictions of highly similar gut communities across colonies of the same termite species and showing clear distinctions between castes within nests. Despite the continued exchange of symbionts via trophallaxis, a subset of bacteria proliferates differentially between soldiers and workers to likely serve different tasks associated with differences in roles and diets. The extremely divergent queen and king gut microbiotas underline the strong influence of division of labour in fungus-growing termites to the symbiotic level and suggests intricate associations between individual termite roles and gut microbiota structure. While our findings provide some indication that worker and soldier guts harbour distinct enzymatic profiles, consistent with differences in diet, the inevitable roles of the non-random assemblies of small sets of dominant royal pair gut symbionts remain to be explored. Our findings set the stage for functional studies that can elucidate the relative importance of bacterial genera across termite castes and the mechanisms underlying shifts in gut compositions, particularly in the royal pair, who initiate their colony with microbial inocula for the first worker cohort, but radically change their gut microbial associations over the course of their extended lifespan.

## Acknowledgements

We thank Z. Wilhelm de Beer, Michael J. Wingfield and the staff and students at FABI, University of Pretoria, for hosting fieldwork, the Oerlemans for permission to sample colonies, and Victoria Challinor, Morten Schiøtt and Justinn Hamilton for comments on a previous draft of the manuscript. The microscopy work was performed at the facilities of the Center for Advanced Bioimaging (CAB) Denmark, University of Copenhagen. This work was supported by a PhD stipend from University of Copenhagen to SO, the EU Horizon 2020 research and innovation program under the Marie Sklodowska-Curie grant agreement No. 660255 to MZ, the British Ecological Society Research grant - Ecologists in Africa (4075-4956) to NAK, a PhD stipend from the CAPES Foundation, Ministry of Education of Brazil, Brasília, Brazil (grant BEX: 13240/13-7) to RRDC, a DFG fellowship (MI 2242/1-1) awarded to AM, and the Villum Kann Rasmussen foundation for a Young Investigator Fellowship (10101) to MP.

## Data deposition

Clean reads are submitted to SRA, GenBank (sample sequences with their accession numbers are under a single SRA submission with the SRA accession: SRP144287).

---

**Supplementary Table S1**. Termite colonies, sampling sites, the number of high-quality 16S rRNA gene sequences obtained from MiSeq amplicon sequencing of the V3-V4 region, and the number of OTUs.

**Supplementary Table S2**. Relative abundances of full bacterial OTUs identified in each termite sterile caste gut sample with full taxonomical levels presented. Click on the (+) sign or the numbers on the left panel to expand the taxon column.

**Supplementary Table S3**. Relative abundances of full bacterial OTUs identified in each termite reproductive caste gut sample with full taxonomical levels presented. Click on the (+) sign or the numbers on the left panel to expand the taxon column.

**Supplementary Table S4**. Termite colonies, sampling sites, sampling year, and castes and ages of termites used for the AZCL experiment, as well as the results of the measurements with one-way PERMANOVA statistical analysis between soldier and worker from different species.

**Supplementary Table S5.** Diversity index values of termite sterile and reproductive caste gut microbiotas, with Mann–Whitney–Wilcoxon statistical analyses of diversity indices between sterile and reproductive castes. Mn: *Macrotermes natalensis*, Od: *Odontotermes* sp., Od.b: *Odontotermes* cf. *badius*, A: *Ancistrotermes*, P: *Pseudacanthotermes*, S: soldier, JS: major soldier, NS: minor soldier, JW: major worker, NW: minor worker, Q: queen, and K: king.

**Supplementary Table S6.** Statistical analyses of Bray-Curtis dissimilarity distances between termite sterile castes, and reproductive caste gut microbiota structures, with analysis of molecular variance (AMOVA) was performed between gut microbiotas using R scripts implemented in mothur between distantly clustered microbial: between the different sterile caste guts from different termite species, and between different royal pair guts from different termite species, also between queen and king guts.

**Supplementary Table S7**. FISH probes used for hybridisations and confocal microscopy.

**Supplementary Table S8**. The results of dada2 analyses of royal pair guts, averaged across three technical replicates (amplifications and sequencing of the same sample). Results are given as relative abundances within samples. The top list provides a comparison of the results to those from the Mothur OTU analyses, with yellow highlights of samples for which the two analyses did not provide the exact same genus-level classification.

**Supplementary Table S9**. The results of dada2 analyses of sterile caste guts, averaged across three technical replicate amplifications and community sequencing. Results are given as relative abundances within samples and labels are: Mn: Macrotermes natalensis, Od: *Odontotermes* sp., Od.b: *Odontotermes* cf. *badius*, S: soldier, JS: major soldier, NS: minor soldier, JW: major worker, NW: minor worker.

**Supplementary Table S10**. Results of linear mixed models, used to assess the main factors that affected OTU Shannon diversity and evenness. Two models were run using the R package vegan, with the colony used as a random effect and the Shannon or evenness indices as response variables. Main effects included in the models were the geographical location (site), the caste and the phylogeny. For the multiple comparisons we performed Tukey post-hoc tests using the lsmeans package in R. Models were constructed using the alpha-diversity indices calculated using the OTU tables from mothur (A), from dada2 (B) or a separate merged analysis using mothur (C).

**Supplementary Table S11**. Results of beta-diversity analysis. For the analyses we used Bray-Curtis distances calculated using the OTU table from the merged mothur analysis of all samples. The OTU table was rarefied at 13000 reads. We used the betapart package in R which calculates the overall beta diversity but also separates beta-diversity in turnover and nestedness. The former refers to beta diversity attributable to species replacement, whereas the latter indicates species loss or gain, that is, richness differences between the samples. We used as main effects, similar to before, the geographical location (site), the caste and the phylogeny, which were evaluated with ANOVA tests. For multiple comparison tests we used Tukey post-hoc tests.

**Supplementary Table S12**. The results of pairwise DESEQ2 comparisons of OTU abundances between castes in *Macrotermes natalensis* and *Odontotermes* spp. Only OTUs with significant differences are given.

**Supplementary Figure S1.**
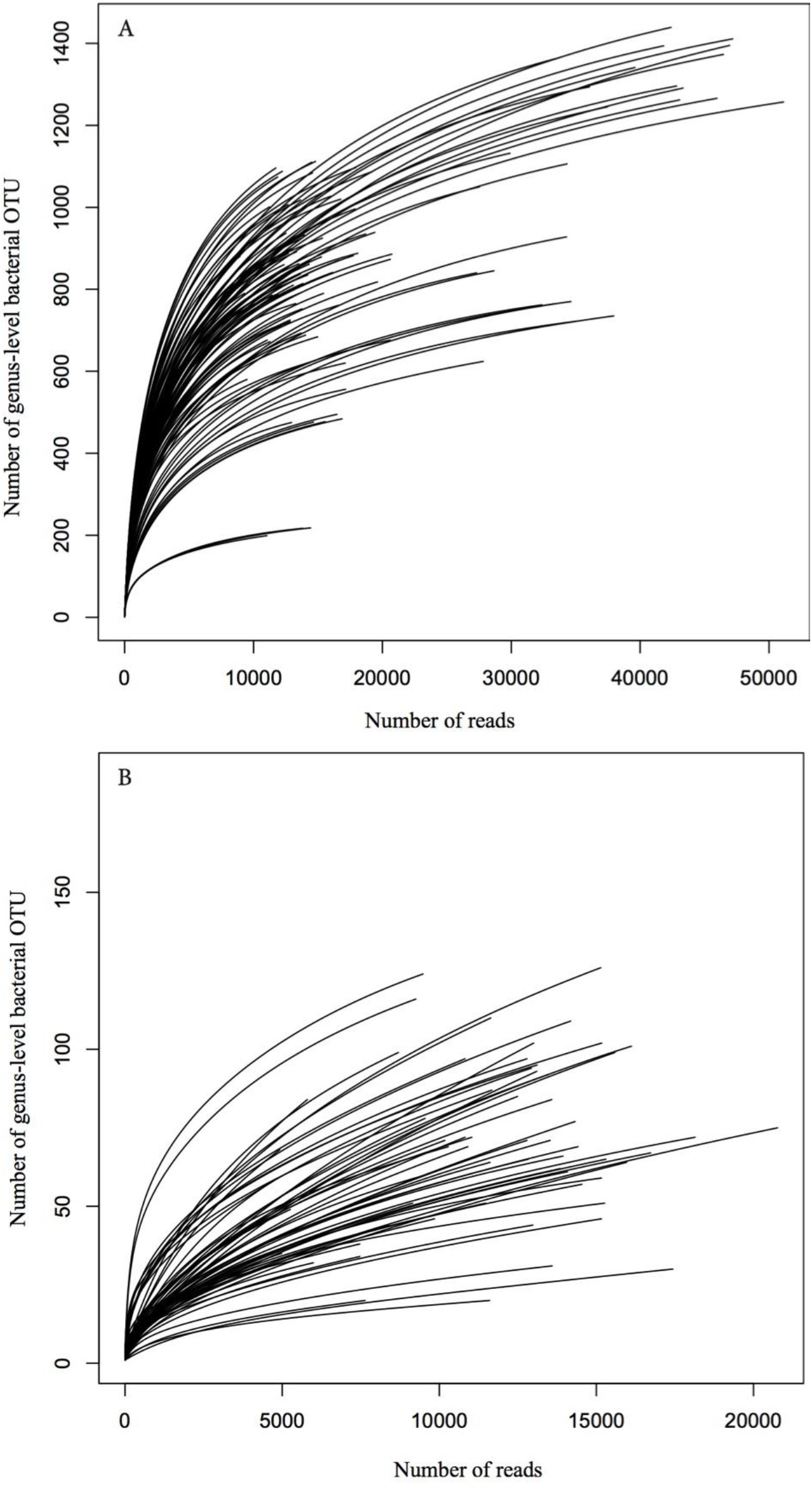
A) Rarefaction curves for sterile castes. B) Rarefaction curves for reproductive castes.

**Supplementary Figure S2.**
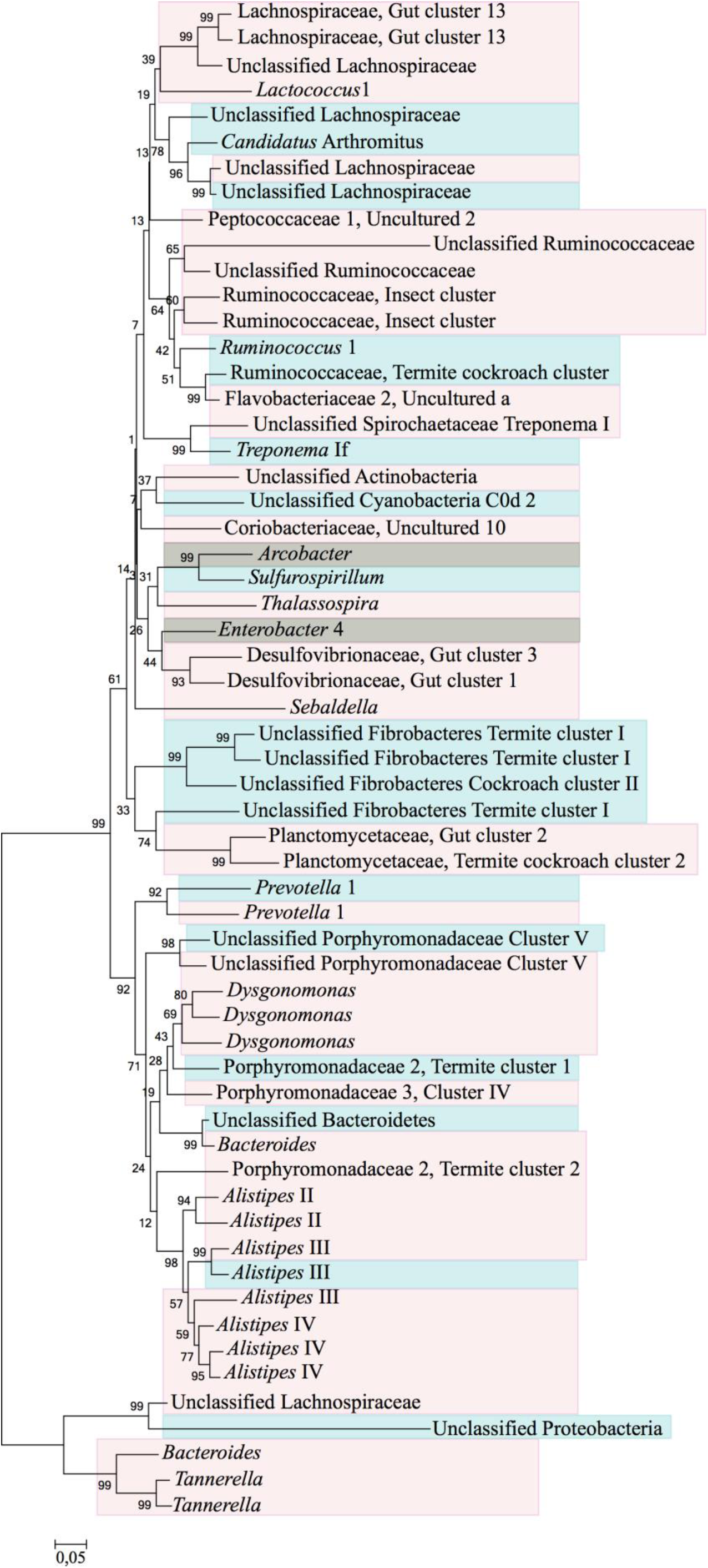
Maximum Likelihood phylogeny of differentially abundant OTUs between sterile castes. Differentially abundant OTUs between *Odontotermes* workers and soldiers highlighted in pink boxes, between *M. natalensis* workers and soldiers in blue boxes, and between workers and soldiers in both termite genera in grey boxes.

**Supplementary Figure S3.**
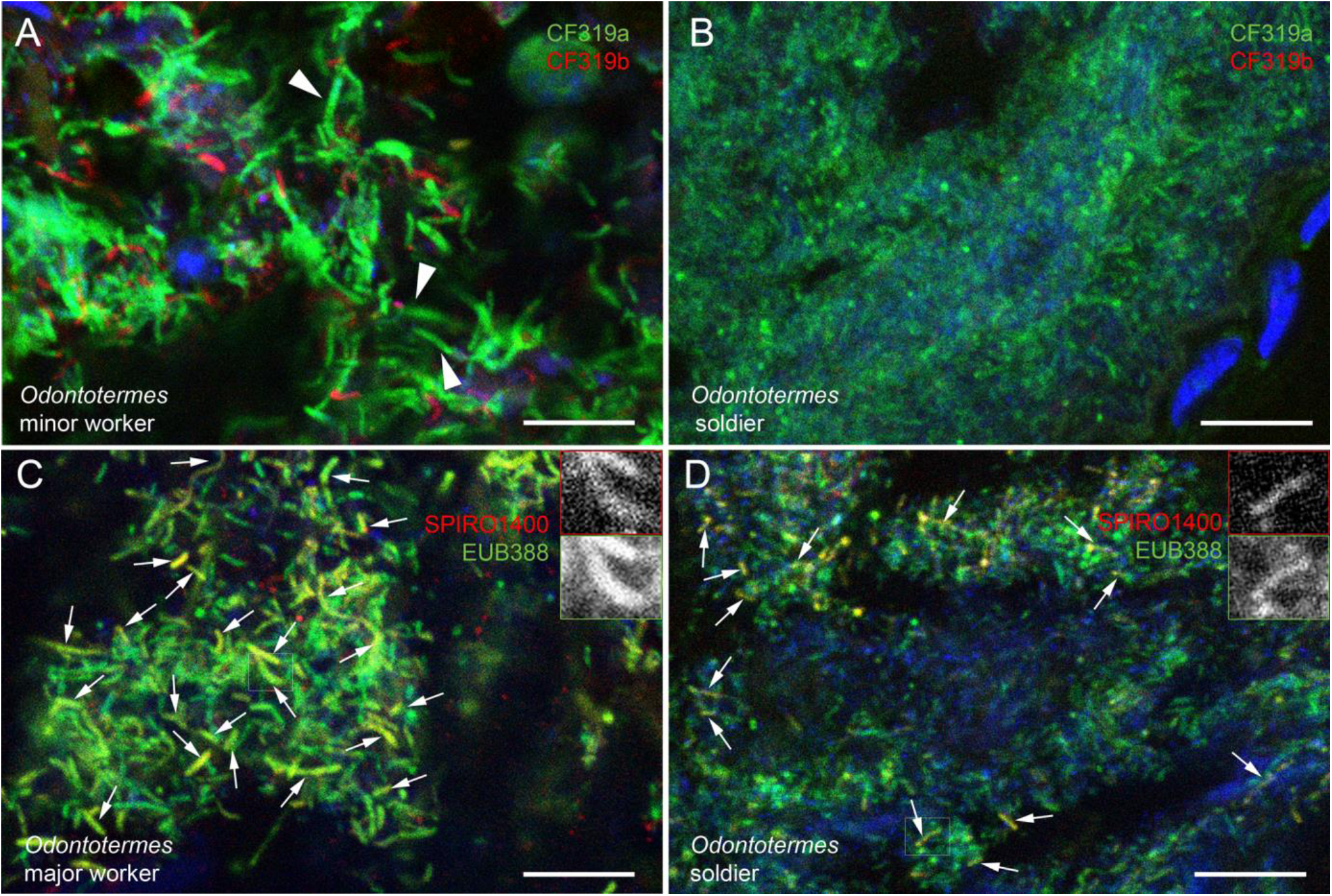
Representative images of fluorescent *in situ* hybridization (FISH) of the remaining three probes (two bacterial taxa) not included in Figure 2G-J. A, B) FISH with two probes (CF319a, green and CF319b, red) targeting members of Bacteroidetes in sterile castes of *Odontotermes* cf. *badius*. Note that bacteria stained with CF319a appear larger in minor workers (arrowheads). C, D) FISH with a probe targeting members of the family Spirochaetaceae in sterile castes of *Odontotermes* cf. *badius*. Scale bars are 10 µm.

**Supplementary Figure S4.**
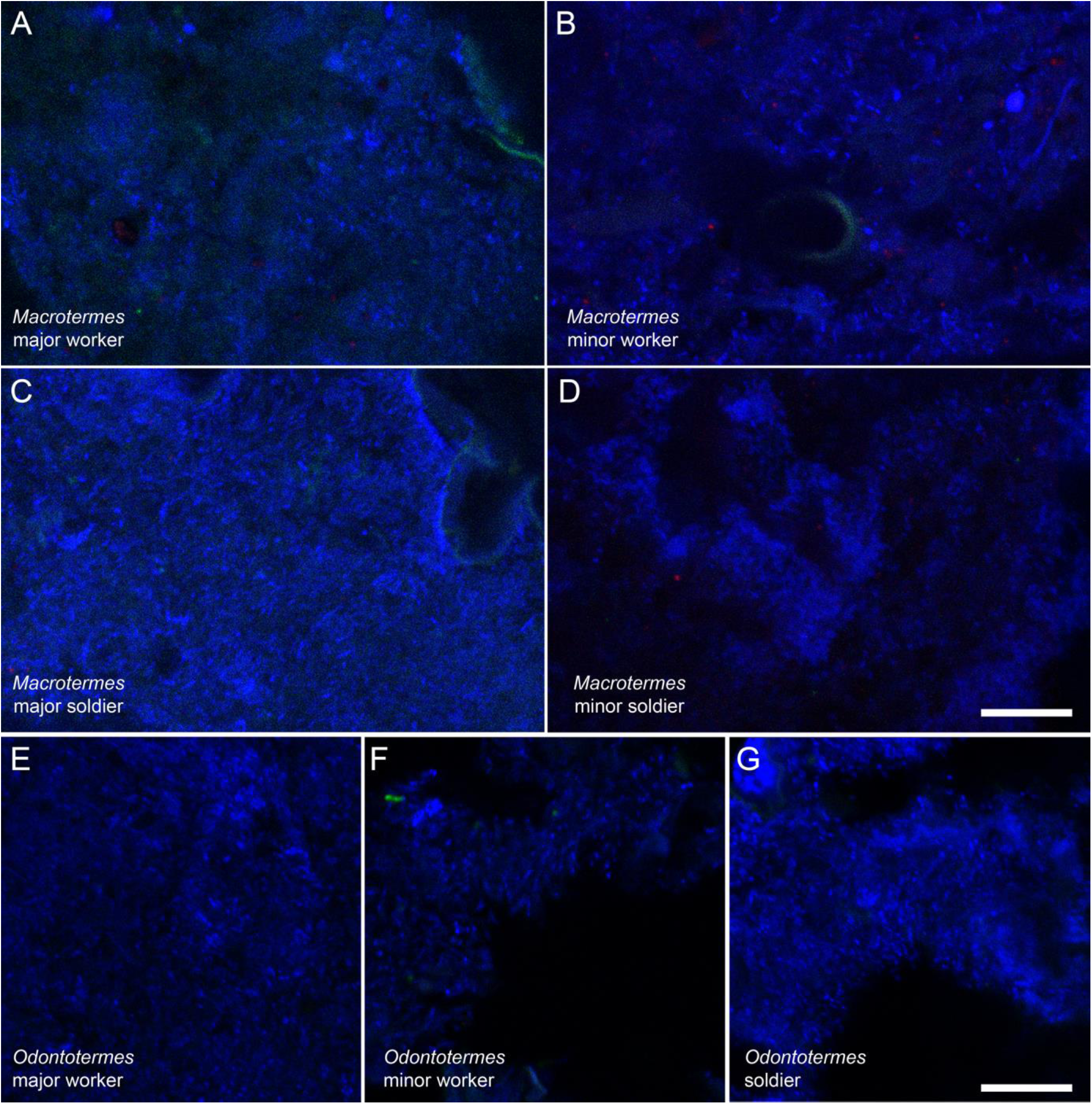
Negative controls of fluorescent *in situ* hybridization using nonEUB388 probe in *M. natalensis* guts (A-D) and *O. badius* guts (E-G). Note the absence of signal. All images are combining three channels: blue (DNA), green (probe staining and autofluorescence), red (autofluorescence). Scale bars are 10 µm.

**Supplementary Figure S5.**
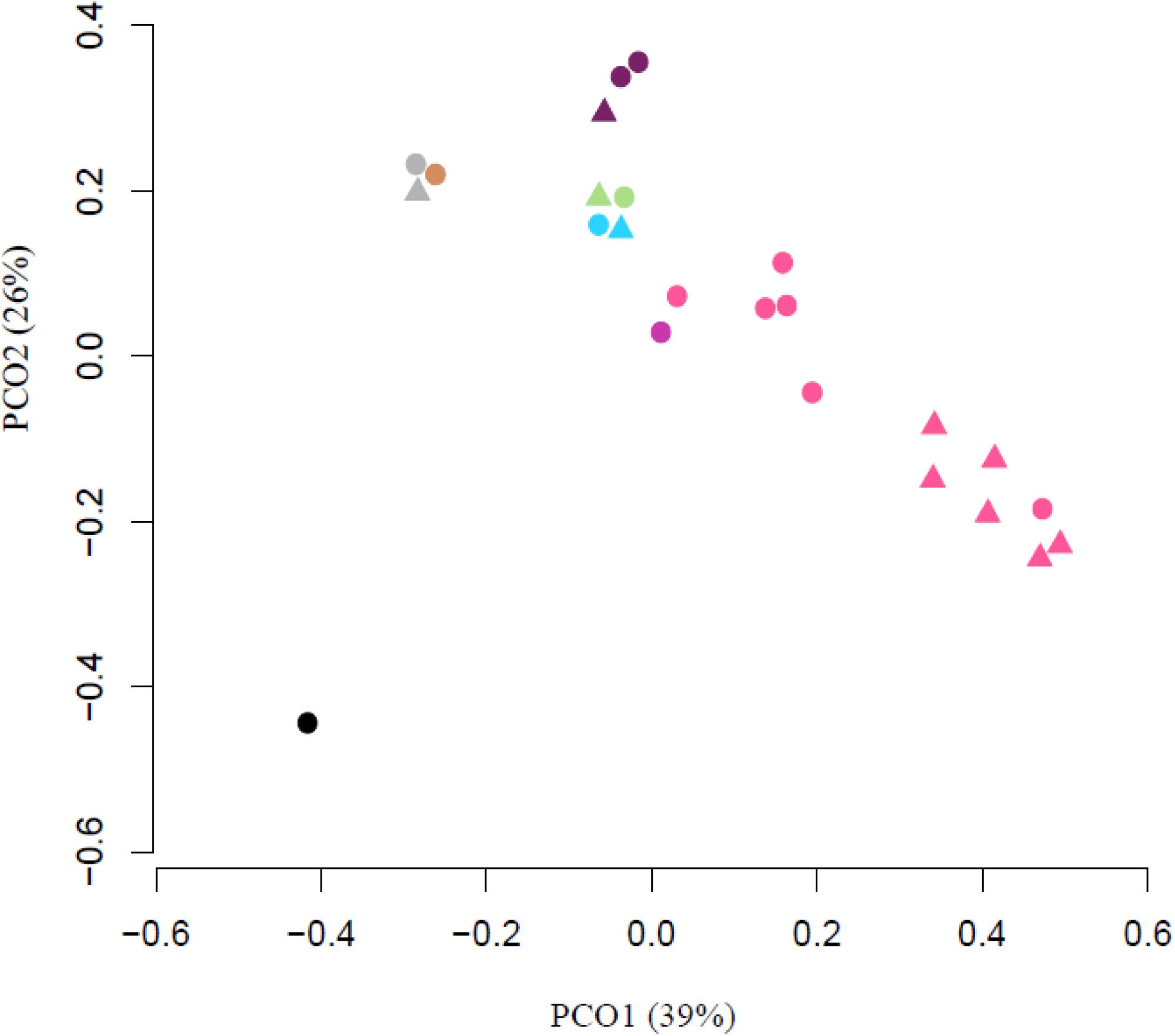
Queen and king gut microbiota similarity analysis (Bray-Curtis) visualised by principal coordinate analysis (PCoA) in R (R core team, 2013). The symbols indicate a royal pair (circle=queen, triangle=king) and colours represents different termite taxa (grey=*Ancistrotermes guineensis*, brown=*Ancistrotermes cavithorax*, green=*Pseudacanthotermes*, blue=*Odontotermes*, light pink=*Macrotermes natalensis*, dark pink=*Macrotermes bellicosus* and purple=*Macrotermes* sp.).

## References

Aanen, D.K., Eggleton, P., Rouland-Lefèvre, C., Guldberg-Frøslev, T., Rosendahl, S., and Boomsma, J.J. (2002) The evolution of fungus-growing termites and their mutualistic fungal symbionts. Proc Natl Acad Sci USA 99: 14887–14892.

Anderson, K.E., Russell, J.A., Moreau, C.S., Kautz, S., Sullam, K.E., Hu, Y.I. et al., (2012) Highly similar microbial communities are shared among related and trophically similar ant species. Mol Ecol 21: 2282–2296.

Badertscher, S., Gerber, C., and Leuthold, R.H. (1983) Polyethism in food supply and processing in termite colonies of *Macrotermes subhyalinus* (Isoptera). Behav Ecol Sociobiol 12: 115–119.

Baselga, A. and Orme, C. D. (2012) betapart: an R package for the study of beta diversity. Methods Ecol Evol 3: 808–812.

Batra, L.R. and Batra, S.W.T. (1979) Termite-fungus mutualism. In: Batra LR (ed). Insect-fungus symbiosis: nutrition, mutualism, and commensalism. Allaheld & Osmun: Montclair. pp 117–163.

Benjamino, J. and Graf, J. (2016) Characterization of the Core and Caste-Specific Microbiota in the Termite, *Reticulitermes flavipes*. Frontiers Microbiol 7: 171.

Berlanga, M., Paster, B.J., Grandcolas, P., and Guerrero, R. (2011) Comparison of the gut microbiota from soldier and worker castes of the termite *Reticulitermes grassei*. Intl Micr 14: 83–93.

Boomsma, J.J. (2009) Lifetime monogamy and the evolution of eusociality. Phil Trans Roy Soc B 364: 3191–3207.

Bourguignon, T., Lo, N., Cameron, S.L., Šobotník, J., Hayashi, Y., Shigenobu, S. et al. (2015) The Evolutionary History of Termites as Inferred from 66 Mitochondrial Genomes. Mol Biol Evol 32: 406–421.

Bourgoignon, T., Lo, N., Dietrich, C., Šobotník, J., Sidek, S., Roisin, Y. et al. (2018) Rampant host switching shaped the termite gut microbiome. Curr Biol 28: 649–654.

Brune, A. and Ohkuma, M. (2011) Role of the Termite Gut Microbiota in Symbiotic Digestion. In: Bignell, D., Roisin, Y., and Lo, N. (eds). Biology of Termites: A Modern Synthesis, 2nd edn. Springer: Dordrecht. pp 439–475.

Brune, A. (2014) Symbiotic digestion of lignocellulose in termite guts. Nature Reviews Microbiology 12: 168–180.

Brune, A. and Dietrich, C. (2015) The Gut Microbiota of Termites: Digesting the Diversity in the Light of Ecology and Evolution. Ann Rev Microbiol 69: 145–166.

Crespi, B.J. and Yanega, D. (1995) The definition of eusociality. Behav Ecol 6: 109–115.

Dietrich, C., Köhler, T., and Brune, A. (2014) The cockroach origin of the termite gut microbiota: patterns in bacterial community structure reflect major evolutionary events. Appl Env Micr 80: 2261–2269.

Donovan, S.E., Eggleton, P., and Bignell, D.E. (2001) Gut content analysis and a new feeding group classification of termites. Ecol Entomol 26: 356–366.

Douglas, A.E. (2009) The microbial dimension in insect nutritional ecology. Functional Ecology 23: 38–47.

Douglas, A.E. (2015) Multiorganismal Insects: Diversity and Function of Resident Microorganisms. Ann Rev Entomol 60: 17–34.

Droge, S., Rachel, R., Radek, R., and Konig, H. (2008) *Treponema isoptericolens* sp. nov., a novel spirochaete from the hindgut of the termite *Incisitermes tabogae*. Intl J Syst Evol Micr 58: 1079– 1083.

Eggleton, P. and Tayasu, I. (2001) Feeding groups, lifetypes and the global ecology of termites. Ecol Res 16: 941–960.

Eggleton, P. (2011) An Introduction to Termites: Biology, Taxonomy and Functional Morphology. In: Bignell, D., Roisin, Y., and Lo, N. (eds). Biology of Termites: A Modern Synthesis. Springer: Dordrecht. pp 349–373.

Ellegaard, K.M., Tamarit, D., Javelind, E., Olofsson, T.C., Andersson, S.G., and Vásquez, A. (2015) Extensive intra-phylotype diversity in lactobacilli and bifidobacteria from the honeybee gut. BMC Genomics 16: 284.

Engel, M.S., Grimaldi, D.A., and Krishna, K. (2009) Termites (Isoptera): Their Phylogeny, Classification, and Rise to Ecological Dominance. American Museum Novitates 3650: 1–27.

Engel, P. and Moran, N.A. (2013) The gut microbiota of insects - diversity in structure and function. FEMS Micr Rev 37: 699–735.

Excoffier, L., Smouse, P.E., and Quattro, J.M. (1992) Analysis of molecular variance inferred from metric distances among DNA haplotypes: application to human mitochondrial DNA restriction data. Genetics 131: 479–491.

Ferguson-Gow, H., Sumner, S., Bourke, A.F.G., and Jones, K.E. (2014) Colony size predicts division of labour in attine ants. Proc R Soc B 281: 20141411.

Gerber, C., Badertscher, S., and Leuthold, R.H. (1988) Polyethism in *Macrotermes bellicosus* (Isoptera). Insectes Soc 35: 226–240.

Graber, J.R. and Breznak, J.A. (2004) Physiology and Nutrition of *Treponema primitia*, an H2/CO2-Acetogenic Spirochete from Termite Hindguts. Appl Env Micr 70: 1307–1314.

Hammer, Ø., Harper, D.A.T., and Ryan, P.D. (2001) PAST: paleontological statistics software package for education and data analysis. Palaentol Elect 31: 139–150.

Hongoh, Y., Ekpornprasit, L., Inoue, T., Moriya, S., Trakulnaleamsai, S., Ohkuma, M. et al. (2006) Intracolony variation of bacterial gut microbiota among castes and ages in the fungus-growing termite *Macrotermes gilvus*. Mol Ecol 15: 505–516.

Hongoh, Y. (2010) Diversity and genomes of uncultured microbial symbionts in the termite gut. Biosci Biotechnol Biochem 74: 1145–1151.

Hongoh, Y. (2011) Toward the functional analysis of uncultivable, symbiotic microorganisms in the termite gut. Cell Mol Life Sci 68: 1311–1325.

Johnson, B.R. (2010) Division of labor in honeybees: form, function, and proximate mechanisms. Behav Ecol Sociobiol 64: 305–316.

Kapheim, K.M., Rao, V.D., Yeoman, C.J., Wilson, B.A., White, B.A., Goldenfeld, N. et al. (2015) Caste-Specific Differences in Hindgut Microbial Communities of Honey Bees (*Apis mellifera*). PLoS ONE 10: e0123911.

Killer, J., Dubna, S., Sedlacek, I., and Svec, P. (2014) *Lactobacillus apis* sp. nov., from the stomach of honeybees (*Apis mellifera*), having an *in vitro* inhibitory effect on the causative agents of American and European foulbrood. Intl J Syst Evol Micr 64: 152–157.

Koch, H. and Schmid-Hempel, P. (2011). Socially transmitted gut microbiota protect bumble bees against an intestinal parasite. Proc Natl Acad Sci USA 108: 19288–19292.

Koenigsknecht, M.J, Theriot, C.M., Bergin, I.L., Schumacher, C.A., Schloss, P.D., and Young, V.B. (2014) Dynamics and Establishment of *Clostridium difficile* Infection in the Murine Gastrointestinal Tract. Inf Immun 83: 934–941.

Koga, R., Tsuchida, T., and Fukatsu, T. (2009) Quenching autofluorescence of insect tissues for *in* situ detection of endosymbionts. Appl Entomol Zool 44: 281–291.

Korb, J. and Hartfelder, K. (2008) Life history and development--a framework for understanding developmental plasticity in lower termites. Biol Rev Camb Phil Soc 83: 295–313.

Kozich, J.J., Westcott, S.L., Baxter, N.T., Highlander, S.K., and Schloss, P.D. (2013) Development of a dual-index sequencing strategy and curation pipeline for analyzing amplicon sequence data on the MiSeq Illumina sequencing platform. Appl Env Micr 79: 5112–5120.

Kwong, W.K., Mancenido, A.L., and Moran, N.A. (2014) Genome Sequences of *Lactobacillus* sp. Strains wkB8 and wkB10, Members of the Firm-5 Clade, from Honey Bee Guts. Genome Announcements 2: e01176–01114.

Kwong, W.K. and Moran, N.A. (2016) Gut microbial communities of social bees. Nature Rev Micr 14: 374–384.

Lanan, M.C., Rodrigues, P.A.P., Agellon, A., Jansma, P., and Wheeler, D.E. (2016) A bacterial filter protects and structures the gut microbiome of an insect. Intl Soc Micr Ecol J 10: 1866–1876.

Leuthold, R.H., Badertscher, S., and Imboden, H. (1989) The inoculation of newly formed fungus comb with *Termitomyces* in *Macrotermes* colonies (Isoptera, Macrotermitinae). Insectes Sociaux 36: 328–338.

Leuthold, R.H., Trietand, H., and Bernd Schildger, B. (2004) Husbandry and breeding of AfricanGiant Termites (*Macrotermes jeanneli*) at Berne Animal Park. Der Zoologische Garten 74: 26–37.

Li, H., Yang, M., Chen, Y., Zhu, N., Lee, C.Y., Wei, J.Q. et al. (2015). Investigation of Age Polyethism in Food Processing of the Fungus-Growing Termite *Odontotermes formosanus* (Blattodea: Termitidae) Using a Laboratory Artificial Rearing System. J Econ Entomol 108: 266– 273

Li, H., Dietrich, C., Zhu, N., Mikaelyan, A., Ma, B., Pi, R. et al. (2016) Age polyethism drives community structure of the bacterial gut microbiota in the fungus-cultivating termite *Odontotermes formosanus*. Environ Micr 18: 1440–1451.

Li, H., Yelle, D.J., Li, C., Yang, M., Ke, J., Zhang, R. et al. (2017) Lignocellulose pretreatment in a fungus-cultivating termite. Proc Natl Acad Sci USA 114: 4709–4714.

Liu, N., Zhang, L., Zhou, H., Zhang, M., Yan, X., Wang, Q. et al. (2013) Metagenomic Insights into Metabolic Capacities of the Gut Microbiota in a Fungus-Cultivating Termite *Odontotermes yunnanensis*. PLoS ONE 8: e69184.

Love, M.I., Huber, W., and Anders, S. (2014) Moderated estimation of fold change and dispersion for RNA-seq data with DESeq2. Genome Biol 15: 550.

Mikaelyan, A., Strassert, J.F., Tokuda, G., and Brune, A. (2014) The fibre-associated cellulolytic bacterial community in the hindgut of wood-feeding higher termites (*Nasutitermes* spp.). Environ Micr 16: 2711–2722.

Mikaelyan, A., Dietrich, C., Köhler, T., Poulsen, M., Sillam-Dussès, D., and Brune, A. (2015a) Diet is the primary determinant of bacterial community structure in the guts of higher termites. Mol Ecol 24: 5284–5295.

Mikaelyan, A., Köhler, T., Lampert, N., Rohland, J., Boga, H., Meuser, K. et al. (2015b) Classifying the bacterial gut microbiota of termites and cockroaches: A curated phylogenetic reference database (DictDb). Syst Appl Micr 38: 472–482.

Mikaelyan, A., Meuser, K., and Brune, A. (2017) Microenvironmental heterogeneity of gut compartments drives bacterial community structure in wood-and humus-feeding higher termites. FEMS Micr Ecol 93: fiw210.

Nalepa, C.A. (2015) Origin of termite eusociality: trophallaxis integrates the social, nutritional, and microbial environments. Ecol Entomol 40: 323–335.

Nobre, T., Rouland-Lefèvre, C., and Aanen, D.K. (2011) Comparative Biology of Fungus Cultivation in Termites and Ants. In: Bignell, D., Roisin, Y., and Lo N (eds). Biology of Termites: A Modern Synthesis, 2nd edn. Springer: Dordrecht. pp 193-210.

Otani, S., Mikaelyan, A., Nobre, T., Hansen, L.H., Koné, N.G.A., Sørensen, S.J. et al. (2014) Identifying the core microbial community in the gut of fungus-growing termites. Mol Ecol 23: 4631–4644.

Otani, S., Hansen, L.H., Sørensen, S.J., and Poulsen, M. (2016) Bacterial communities in termite fungus combs are comprised of consistent gut deposits and contributions from the environment. Micr Ecol 71: 207–220.

Paulson, J.N., Stine, O.C., Bravo, H.C., and Pop, M. (2013) Differential abundance analysis for microbial marker-gene surveys. Nature Meth 10: 1200–1202.

Pinheiro, J., Bates, D., DebRoy, S., Sarkar, D., and R Core Team (2018). nlme: Linear and Nonlinear Mixed Effects Models. R package version 3.1-137, https://CRAN.R-project.org/package=nlme.

Poulsen, M., Hu, H., Li, C., Chen, Z., Xu, L., Otani, S. et al. (2014) Complementary symbiont contributions to plant decomposition in a fungus-farming termite. Proc Natl Acad Sci USA 111: 14500–14505.

R Core Team (2013). R: A language and environment for statistical computing. R Foundation for Statistical Computing, Vienna, Austria. URL http://www.R-project.org/.

Rahman, A.N., Parks, D.H., Willner, D.L., Engelbrektson, A.L., Goffredi, S.K., Warnecke, F. et al. (2015) A molecular survey of Australian and North American termite genera indicates that vertical inheritance is the primary force shaping termite gut microbiomes. Microbiome 3: 5.

ouland-Lefèvre, C., Inoue, T., and Johjima, T. (2006) *Termitomyces*. In: König, H. and Varma, A. (eds). Intestinal Microorganisms of Soil Invertebrates. Springer-Verlag: Berlin Heidelberg. pp 335-350.

Roy, V., Girondot, M., and Harry, M. (2015) The Distribution of *Wolbachia* in *Cubitermes* (Termitidae, Termitinae) Castes and Colonies: A Modelling Approach. PLoS ONE 10: e0116070.

Russel, J., Thorsen, J., Brejnrod, A.D., Bisgaard, H., Sørensen, S.J., and Burmølle, M. (2018). DAtest: a framework for choosing differential abundance or expression method. bioRxiv preprint first posted online Jan. 2, 2018; doi: http://dx.doi.org/10.1101/241802.

Salunke, B.K., Salunkhe, R.C., Dhotre, D.P., Khandagale, A.B., Walujkar, S.A., Kirwale, G.S. et al. (2010) Diversity of Wolbachia in Odontotermes spp. (Termitidae) and Coptotermes heimi (Rhinotermitidae) using the multigene approach. FEMS Microbiology Letters 307: 55–64.

Sands, W.A. (1960) The initiation of fungus comb construction in laboratory colonies of *Ancistrotermes guineensis* (Silvestri). Insectes Soc 7: 251–263.

Sands, W.A. (1969) The association of termites and fungi. In: Krishna, A. and Weesner, F. (eds). Biology of Termites. Academic Press: New York. pp 495-524.

Sapountzis, P., Zhukova, M., Hansen, L.H., Sørensen, S.J., Schiøtt, M., and Boomsma, J.J. (2015) *Acromyrmex* Leaf-Cutting Ants Have Simple Gut Microbiota with Nitrogen-Fixing Potential. Appl Env Micr 81: 5527–5537.

Schloss, P.D., Westcott, S.L., Ryabin, T., Hall, J.R., Hartmann, M., Hollister, E.B. et al. (2009) Introducing mothur: Open-Source, Platform-Independent, Community-Supported Software for Describing and Comparing Microbial Communities. Appl Env Micr 75: 7537–7541.

Sharon, G., Segal, D., Ringo, J.M., Hefetz, A., Zilber-Rosenberg, I., and Rosenberg, E. (2010) Commensal bacteria play a role in mating preference of Drosophila melanogaster. Proc Natl Acad Sci USA 107: 20051–20056.

Tamura, K., Dudley, J., Nei, M., and Kumar, S. (2007) MEGA4: Molecular Evolutionary Genetics Analysis (MEGA) software version 4.0. Mol Biol Evol 24: 1596–1599.

Tarpy, D.R., Mattila, H.R., and Newton, I.L.G. (2015) Development of the Honey Bee Gut Microbiome throughout the Queen-Rearing Process. Appl Env Micr 81: 3182–3191.

Turnbaugh, P.J., Ley, R.E., Mahowald, M.A., Magrini, V., Mardis, E.R., and Gordon, J.I. (2006) An obesity-associated gut microbiome with increased capacity for energy harvest. Nature 444: 1027–1131.

Veivers, P.C., Mühlemann, R., Slaytor, M., Leuthold, R.H., and Bignell, D.E. (1991) Digestion, diet and polyethism in two fungus-growing termites: *Macrotermes subhyalinus* Rambur and *M. michaelseni* Sjøstedt. J Insect Phys 37: 675–682.

Watanabe, H. and Tokuda, G. (2010) Cellulolytic Systems in Insects. Ann Rev Entomol 55: 609– 632.

## Supplementary Information References

Amann, R.I., Binder, B.J., Olson, R.J., Chisholm, S.W., Devereux, R., and Stahl, D.A. (1990) Combination of 16S rRNA-targeted oligonucleotide probes with flow cytometry for analyzing mixed microbial populations. Appl Env Micr 56: 1919–1925.

Daly, K. and Shirazi-Beechey, S.P. (2003) Design and evaluation of group-specific oligonucleotide probes for quantitative analysis of intestinal ecosystems: Their application to assessment of equine colonic microflora. FEMS Micr Ecol 44: 243–252.

Hallberg, K.B., Coupland, K., Kimura, S., and Johnson, D.B. (2006) Macroscopic streamer growths in acidic, metal-rich mine waters in North Wales consist of novel and remarkably simple bacterial communities. Appl Env Micr 72: 2022–2030.

Manz, W., Amann, R., Ludwig, W., Vancanneyt, M., and Schleiferl, K.-H. (1996) Application of a suite of 16s rRNA-specific oligonucleotide probes designed to investigate bacteria of the phylum cytophaga-f lavobacter-bacteroides in the natural environment. Microbiology 142: 1097–1101.

Rigottier-Gois, L., Rochet, V., Garrec, N., Suau, A., Doré, J. (2003) Enumeration of *Bacteroides* species in human faeces by fluorescent in situ hybridisation combined with flow cytometry using 16S rRNA probes. Syst Appl Micr 26: 110–118.

